# PPR767 affects plant architecture and drought resistance by modulating complex I activity and ROS content in rice

**DOI:** 10.1101/2024.07.17.603968

**Authors:** Leilei Peng, Haijun Xiao, Yanghong Xu, Zhihao Huang, Xuan Yang, Chen Lv, Linghui Huang, Jun Hu

## Abstract

The RNA-binding proteins (RBPs) encoded by nucleus are essential for the metabolism of RNAs in eukaryotes. The pentatricopeptide repeat (PPR) proteins, a large subset of RBPs, participate in organellar RNA processing for plant development and reproduction. Here, we identified an E-type PPR protein, PPR767, which functions in mitochondria. Knocking out *PPR767* resulted in shorter plant height, thinner stems, shorter and narrower blades, and consequently affected yield traits, compared to those of the wild type. PPR767 primarily participated in the editing of 4 sites, nad1-674, *nad3*-155, *nad3*-172, and *nad7*-317. And PPR767 interplayed with MORF1 and MORF8, suggesting the editosome in rice is complicated. Meanwhile, the activity of mitochondrial complex I was decreased, and the structure of mitochondria was compromised in the mutants. Furthermore, mutation of *PPR767* influenced rice drought tolerance and the expression levels of genes involved in the reactive oxygen species (ROS) accumulation. These findings suggest that *PPR767* guarantees the complex I activity by properly regulating the RNA editing efficiency of mitochondrial genes and affects drought tolerance by modulating ROS content in rice, providing valuable insights into the mechanisms by which PPRs fulfil their functions.

## Introduction

As a semiautonomous organelle, it is generally accepted that mitochondria originate from an eubacterial ancestor, ɑ-proteobacteria, which is embraced by eukaryotic cells through endosymbiosis (Gray et al., 2001). Within the host cell, mitochondria generate the energy through respiration (Knoop, 2004). During descending, the majority of eubacterial genes are either lost or migrate into the host genome (Brandvain and Wade, 2009). In land plants, mtDNAs comprise approximately 60-70 genes, the majority of which are related to the production of various subunits of respiratory complexes I (NADH dehydrogenase), III (cytochrome *c* reductase or *bc_1_*), IV (cytochrome *c* oxidase), and V (also denoted as subunits of ATP-synthase), as well as cytochrome *c* biogenesis (CCM) factors essential for electron transport and mitochondrial function (Small et al., 2020). The maturation of mtRNAs from primary transcripts of mtDNAs involves multiple complex post-transcriptional processes, including endonucleolytic cleavage, maturation of both 5’ and 3’ termini, splicing of group I and/or Ⅱ intron sequences, and RNA editing (Hammani and Giege, 2014; Binder et al., 2016).

RNA editing is a posttranscriptional process that challenges the central dogma rule by producing RNAs that differ from their template genome sequences (Knoop, 2011). The phenomenon of RNA editing was initially observed in 1986, the transcript of cytochrome *c* oxidase subunit 2 (*cox2*) from *Trypanosoma brucei* mitochondria contained 4 nucleotides that were not encoded in DNA (Benne et al., 1986). In flowering plants, RNA editing was first reported in 1989 by three independent research groups in wheat and *Oenothera* (Covello and Gray, 1989; Gualberto et al., 1989; Hiesel et al., 1989). Massive studies have identified several types of RNA editing, including the conversion of cytidines (C) to uridine (U), the insertions and deletions of uridine (U), the deamination of adenosine (A) to inosine (I), and the insertion of guanosine (G) (Chen et al., 1987; Thomas et al., 1988; Benne, 1989; Walkley and Li, 2017). In plants, RNA editing occurs in both chloroplasts and mitochondria, with the majority being is C-to-U substitution (Knie et al., 2016). Compared to the 30-40 editing events in the chloroplasts of land plants, mitochondria likely possess more editing events, numbering in hundreds (Bentolila et al., 2012; Xiao et al., 2018a). Although C-to-U events alter the genetic information encoded by DNA, the resulting amino acids tend to be highly conserved throughout the evolution of terrestrial plants, suggesting a conserved mechanism to compensate the mutations of organellar DNA (Gualberto et al., 1989). The occurrence of C-to-U RNA editing requires nucleus-encoded RNA binding proteins (RBPs) or cofactors to form the editosome and execute the deaminase reaction (Barkan and Small, 2014). Among these RBPs, pentatricopeptide repeat proteins (PPRs) have been identified as a class of key components of the editosome with lots of scientific evidence (Kotera et al., 2005).

PPR proteins are typically characterized by a tandem array of approximately 35 degenerate amino acid repeat motifs (Small and Peeters, 2000). PPR motifs are divided into three types: P motif (canonical 35 amino acid PPR motif), L motif (35-36 amino acids), and S motif (31 amino acids) (Barkan and Small, 2014). PPRs are grouped into two subfamilies: the P subfamily, which is identified by containing only the P motif, and the PLS subfamily, which contains additional types of motifs (Lurin et al., 2004). PLS subfamily PPR proteins are further divided into E, E+, and DYW subgroups on the basis of their C-terminal domains (Lurin et al., 2004; Cheng et al., 2016). Generally, PPRs participate in RNA metabolism via recognizing and binding to target RNAs. The crystal structure of PPR proteins has shown that each PPR motif forms a hairpin of ɑ-helices, including helix a and helix b (Yin et al., 2013). Structure analysis revealed that the residues within the first ɑ helix (position 6) and at the end of the loop region (position 1’) are responsible for RNA sequence recognition (Ke et al., 2013; Yin et al., 2013; Hall, 2016). This phenomenon is generally consistent with a previously proposed recognition mode in bioinformatic analysis (Takenaka et al., 2013). However, different types of PPRs are involved in different RNA processing pathways. P-type PPR proteins are involved in RNA splicing, editing, cleavage, stability, transcription, and translation of target RNAs (Chateigner-Boutin et al., 2008; Schmitz-Linneweber and Small, 2008; Barkan and Small, 2014; Rugen et al., 2019; Waltz et al., 2019). PLS subfamily PPR proteins primarily participate in C-to-U RNA editing (Schmitz-Linneweber and Small, 2008; Barkan and Small, 2014). The nature of C-to-U RNA editing is a deamination reaction, with deaminase being the key enzyme involved in the reaction. Several reports have confirmed that the DYW domain of PPRs possesses deaminase activity, implying that non-DYW-type PPR proteins lacking this function may recruit DYW proteins or other cofactors for deamination (Salone et al., 2007; Diaz et al., 2017; Oldenkott et al., 2019). Mutations of PPR proteins affect the metabolism of target RNAs and influence organelle complex activity, resulting in a series of phenotypes (Chateigner-Boutin et al., 2008; Kim et al., 2009; Xiao et al., 2018a). In chloroplast, mutation of OsPGL1 results in defective photosynthetic complex and causes pale green leaves, and the PPR protein RARE1 affects abiotic stress resistance by mediating the editing of *accD* transcripts in *Arabidopsis* (Xiao et al., 2018a; Huang et al., 2023). In mitochondria, knockout of AtRTP7 impacts the ROS content and plant immunity by regulating complex I activity (Yang et al., 2022b). In rice, PPS1 and PPR756, both of which affect pollen sterility, as well as SOP10 which is related to cold resistance, showed reduced complex I activity after mutation in rice (Xiao et al., 2018b; Zhang et al., 2020; Zu et al., 2023).

In addition to PPR proteins, several other nucleus-encoded cofactors are involved in RNA editing as constituents of the editosome, including multiple organellar RNA editing factors (MORFs) or RNA editing interacting proteins (RIPs), organelle RNA recognition motif (ORRM), protoporphyrinogen IX oxidase 1 (PPO1), and organelle zinc fingers (OZs) (Takenaka et al., 2012; Sun et al., 2013; Zhang et al., 2014; Sun et al., 2015). Based on previous reports, MORFs may recruit DYW-type PPRs for other PPRs without deaminase activity in editing events (Guillaumot et al., 2017; Bentolila et al., 2021; Yang et al., 2022c). The ORRMs identified in Arabidopsis and maize are associated with the editing of several sites (Sun et al., 2013; Shi et al., 2016). It has been shown that PPO1 interacts with MORF proteins, and mutations in PPO1 result in editing defects at several sites (Zhang et al., 2014). Moreover, OZ1 was also characterized as a co-factor of an RNA editosome by interacting with ORRM1 and MORFs in chloroplasts (Sun et al., 2015). OZ2 is involved in intron splicing of both several *nad* genes and *rps3* in Arabidopsis mitochondria and is also related to the activity of respiratory complex I (Bentolila et al., 2021).

In this study, we identified a mitochondrion-targeted E-type PPR protein, PPR767, in rice. As an RBP, PPR767 directly binds to and participates in 4 RNA editing sites of 3 mitochondrial genes, *nad1*, *nad3*, and *nad7*. The results confirmed the interaction between PPR767 with both MORF1 and MORF8, suggesting a complicated editosome in mitochondria. Interestingly, all of the target genes are subunits of complex I, the activity of which is obviously reduced in *ppr767* mutants. Moreover, the mitochondria in mutants lacks distinctive crista structure and clear inner membrane systems. Meanwhile, the ROS content is dramatically elevated under drought treatment in the mutants than in the wild type. In conclusion, PPR767 is involved in RNA editing by interacting with MORFs to ensure the activity of complex I and impact plant development and drought tolerance in rice.

## RESULTS

### The *PPR767* mutants show a growth reduction phenotype

To gain insight into the functions of PPR proteins in rice, we identified three mutants of *PPR767* (*LOC_Os05g24150*), named as *ppr767-1*, *ppr767-2*, and *ppr767-3,* with the ZH11 (*O. sativa subsp. japonica*) background were selected for this study. The mutants that exhibited 1 bp insertion or deletion in the first exon of *PPR767* were identified, and all of these mutations resulted in a premature stop codon (Figure **1A**). We also constructed the transgenic vector pCAMBIA2300 containing *PPR767*, driven by its native promoter, to rescue the phenotypes of three mutants, respectively, named as *ppr767-1*-C, *ppr767-2*-C, and *ppr767-3*-C. Real-time quantitative PCR (RT-qPCR) was carried out to verify the expression levels of *PPR767*. The results suggested that the expression levels of *PPR767* were significantly decreased in the three knockout (KO) lines, while were restored in the complemented plants (Figure **1B**).

**Figure 1.**
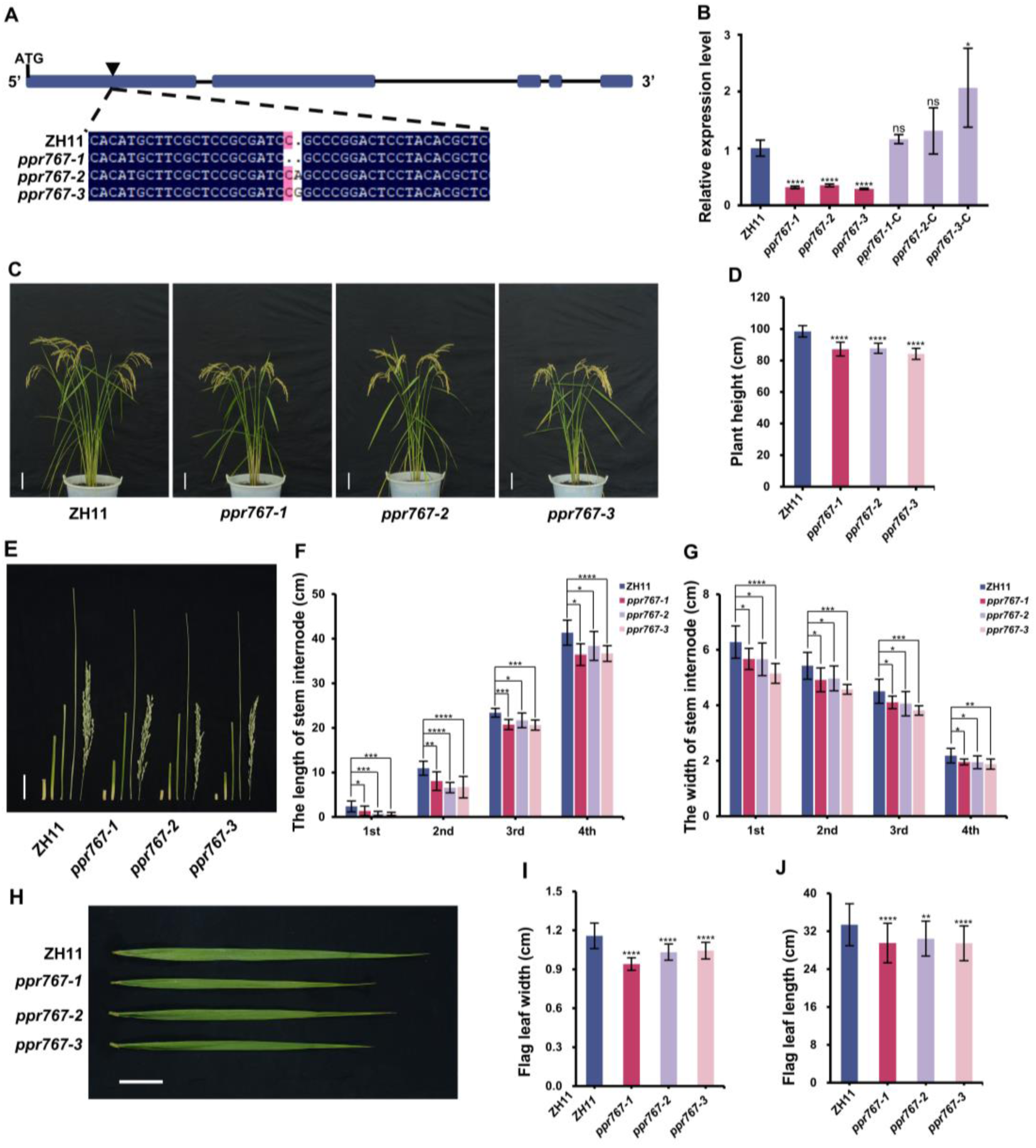
Phenotypic comparisons between the wild type (ZH11) and the mutants. **A**, Schematic view of the mutation site of the *PPR767* gene and alignment of sequences of the wild type and mutated alleles generated by CRISPR-Cas9. The deep blue box and black straight line indicate exons and introns respectively. **B**, Expression analysis of *PPR767* in ZH11 and related transgenic lines (knockout and complemented lines) by real-time quantitative RT-PCR (RT-qPCR). Data are means ± SDs (standard deviations) of three biological replicates. **C**, Gross morphology of ZH11 and the mutants at the mature stage. Scale bar = 10 cm. **D**, Statistical data for plant height at the mature stage. Data are means ± SDs (n = 50). **E**, Comparison of internodes of main tiller between the wild type and three mutant lines. Scale bar = 5 cm. **F** to **G**, Statistical data for the length of stem internodes **(F)**, and width of stem internodes **(G)**, respectively. Data are means ± SDs (n = 10). **H**, Flag leaf comparison between the wild type and three mutant lines. Scale bar = 5 cm. **I** to **J**, Statistical data for flag leaf width **(I)** and flag leaf length **(J)** of ZH11 and three mutant lines. Data are means ± SDs (n = 50). Asterisks indicate significant differences by Student’s two-tailed *t* test (*\*p* < 0.05, *\*\*p* < 0.01, *\*\*\*p* < 0.005, *\*\*\*\*p* < 0.001, ns, no significance).

Compared to the wild type (WT), the plants of the mutants were shorter at the mature stage (Figure **1C-D**). Further investigation showed that this phenotype existed stably at seedling stage, indicating that this characteristic persisted throughout the entire growth period (Supplemental Figure S**1A-B**). To figure out the reason causes this phenotype, the lengths of the internodes of the main tiller were measured at the mature stage. The results revealed that, compared to the WT, the length of every internode was significantly decreased in the mutants (Figure **1E-F**). Meanwhile, the widths of all four stem internodes of the mutants were significantly thinner than those of the WT (Figure **1G**). Furthermore, both the length and width of the flag leaves on the main tiller of the mutants at the mature stage were significantly lower compared to those in WT, suggesting that PPR767 is important for growth duration in rice (Figure **1H-J**).

To observe these alterations at the histological level, histological section analysis on both the internodes and leaves at the filling stage was implemented. Transections of the stems further verified that all the four internodes of three mutants were thinner than those of the WT respectively (Figure **2A**). For stem width, the main determinants are the number, size and interval of vascular bundles. Statistical analysis of the number of adaxial vascular bundles (ADVs) or abaxial vascular bundles (ABVs) showed the number of both ADVs and ABVs were significantly decreased in the mutants than in the wild type (Figure **2B-D**). We also measured the size of ABVs and ADVs, however, compared to WT, the mutants of *PPR767* showed no significant difference in either (Supplemental Figure **S2A-B**). Meanwhile, the interval of ADVs was measured, and the results showed that it was reduced in the mutants (Figure **2E**). These findings indicate that the changes in stem thickness primarily result from the decreased number of both ADVs and ABVs and the reduced interval between ADVs. In addition, we investigated the reasons for the narrower leaves of the mutants. Statistical analysis showed the number of SVs significantly decreased in the mutants, while no significant difference about the number of LVs was detected (Figure **2F-G**, Supplemental Figure **S2C**). In addition to the number of SVs and LVs, the size of the midveins, LVs, and SVs also influence leaf width. However, no remarkably difference were detected between the WT and the mutants (Supplemental Figure **S2D-F**). In addition, the interval of SVs is one of the major factors that impacts the width of leaf. Research indicated that the interval of SVs was minished in the mutants (Figure **2H**). These results demonstrate that the altered leaf width is primarily due to fewer SVs and their reduced intervals.

**Figure 2.**
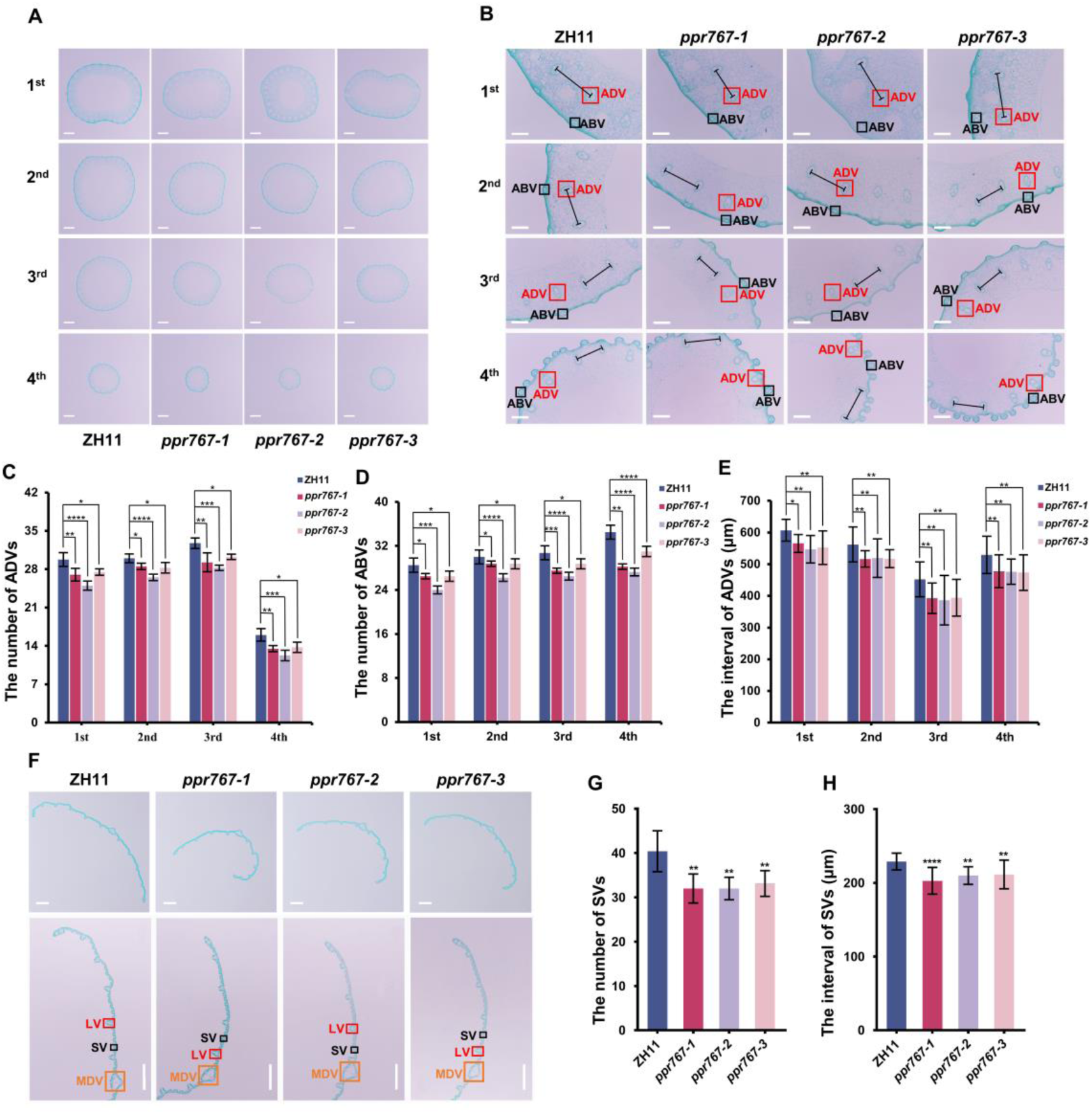
Comparison about cross-sections of internodes and leaves between ZH11 and the mutants. **A**, Cross-sections of each internode of ZH11 and three mutants. 1st, 2nd, 3rd, and 4th indicate the first, second, third and fourth internode counts from bottom to top, respectively. Scale bar = 1000 μm. **B**, Magnification of cross-sections of each internode from ZH11 and the mutants. 1st, 2nd, 3rd, and 4th indicate the first, second, third and fourth internode counts from bottom to top respectively. Scale bar = 250 μm. Red and black boxes indicate the adaxial vascular bundle (ADV) and abaxial vascular bundle (ABV) respectively. The black lines with ends indicate intervals between ADVs. **C** to **E**, Statistical data for the number of ADVs ((**G**), n = 5), the number of ABVs ((**D**), n = 5), and the interval of ADVs ((**E**), n = 20). Data are means ± SDs. **F**, Cross-sections and corresponding magnification about flag leaves of ZH11 and the mutants. Scale bar =1000 μm. The red, black and orange boxes indicate the large vascular bundle (LV), small vascular bundle (SV) and midvein (MDV), respectively. **G** to **H**, Statistical data for the number of SVs ((**G**), n = 5), and the interval of SVs ((**H**), n = 20) of flag leaf cross-sections from ZH11 and the mutants. Data are mean ± SDs. Asterisks indicate significant differences by Student’s two-tailed *t* test (*\*p* < 0.05, *\*\*p* < 0.01, *\*\*\*p* < 0.005, *\*\*\*\*p* < 0.001, ns, no significance).

In order to investigate whether the mutation of *PPR767* influenced the rice yield, we measured the number of tillers, primary branch, secondary branch, and grains per panicle. Statistical analysis showed that, compared to the WT, both the number of primary branch and grains per panicle were reduced in the mutants, whereas the number of tillers and secondary branches were not significantly influenced (Supplemental Figure **S3A-D**). These results indicate that the mutation of *PPR767* may reduce the rice yield.

### PPR767 is a mitochondria-targeted E-type PPR protein

Motif prediction analysis by the PPR (https://ppr.plantenergy.uwa.edu.au/) website and NCBI BLAST (https://blast.ncbi.nlm.nih.gov/Blast.cg) revealed that *PPR767* encodes a putative E-type PPR protein comprising 767 amino acids, containing 13 PLS motifs (including 4 P motifs, 5 L motifs, and 4 S motifs) and one each of the E and E+ domains at C-terminus (Figure **3A**). RT-qPCR revealed that *PPR767* was ubiquitously expressed in all tissues, including seedling, panicle, anther, leaf, root, and stem (Figure **3B**).

**Figure 3.**
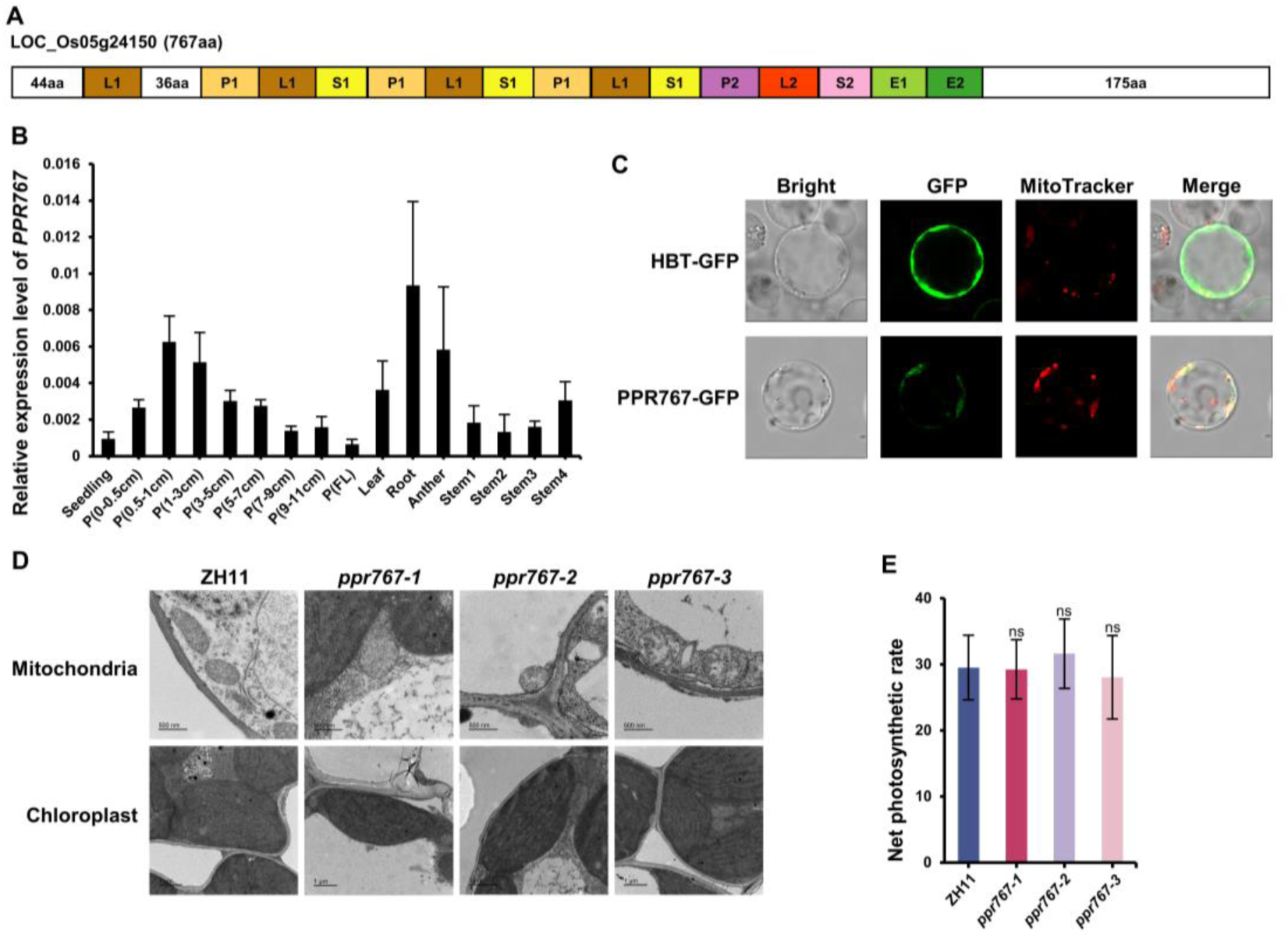
PPR767 is an E-type PPR protein that is located in mitochondria. **A**, Schematic diagram of the protein domain structure. P, L, S, and E indicate P, L, S, and E motif respectively. **B**, RT-qPCR analysis of the expression levels of *PPR767* in different tissues. P, panicle; FL, full length; stem1/2/3/4, the stem internode from bottom to top. Data are means ± SDs of three biological replicates. **C**, Subcellular localization of PPR767. GFP, green fluorescent protein; HBT-GFP, empty vector; MitoTracker Red, mitochondrial red fluorescent dye. **D**, Transmission electron microscopy (TEM) images show the ultrastructure of mitochondria and chloroplast of ZH11 and the mutants. **E**, Comparison of the net photosynthetic rate of ZH11 and the mutants. Data are means ± SDs (n = 8). ns, no significance.

To determine the subcellular localization of PPR767, the CDS of *PPR767* was fused with green fluorescent protein (GFP). The GFP fluorescence of the fusion protein was observed with confocal microscopy and co-localized with the fluorescent signal of MitoTracker Deep Red, which specifically stains mitochondria (Figure **3C**). This result indicates that PPR767 functions within mitochondria. Meanwhile, in order to further verify this conclusion, the structure of mitochondria and chloroplasts was observed via transmission electron microscopy (TEM). The results showed that the chloroplasts exhibited an extensive and normal thylakoid membrane system in both the WT and the mutants, while in contrast to the normal fusiform-shaped mitochondria with a well-regulated and constant crista structure in the WT, the mitochondria of the mutants lacked distinctive crista structure and clear inner membrane systems (Figure **3D**). These alterations in mitochondrial structure in the mutants indicate that the mutation of *PPR767* specifically impacts mitochondrial development and further support the notion that PPR767 functions in mitochondria. In addition, we measured the net photosynthetic rate at the midpoint of flag leaves from ZH11 and three mutants at seven days after heading using the LI6800 system. The net photosynthetic rate of the mutants did not significantly differ from that of ZH11, which further suggested that PPR767 was not associated with the development or function of chloroplasts (Figure **3E**). Based on the aforementioned results, we presume that PPR767 is localized to and executes its function within mitochondria.

### PPR767 participates in the editing of 4 sites in 3 mitochondrial genes

In light of prior research, PPR767, a PPR protein, was hypothesized to function as an RNA-binding protein (RBP), engaging in posttranscriptional modification of one or more RNAs encoded by mitochondrial genes. In order to confirm this hypothesis, total RNA was extracted from the leaves of ZH11 and three mutants at the same developmental stage. All recognized editing sites within the mitochondrial and chloroplast genes were amplified via reverse transcription PCR (RT-PCR), and the amplicons were sequenced via Bulk Sanger sequencing assay. The results showed that no editing sites within the chloroplast genes exhibited significantly altered editing efficiency in the mutants. However, in comparison to the WT, the editing efficiency of 4 sites (*nad1*-674, *nad3*-155, *nad3*-172, and *nad7*-317) within 3 mitochondrial genes was altered in the mutants (Figure **4A**). These results were in accordance with the conclusion that PPR767 is localized to mitochondria. Further investigation revealed that the editing efficiency of *nad1*-674 and *nad3*-155 was decreased in the mutants, preventing the conversion of codon UCU (Ser) to UUU (Phe) and of codon CCG (Pro) to CUG (Leu), respectively (Figure **4A**). Moreover, the editing efficiency of *nad3*-172 was also diminished in the mutants, leading to CUA (Leu) not be converted to UUA (Leu), however, the message of amino acid (aa) remained unchanged due to the degeneracy property of the codon (Figure **4A**). Notably, the editing efficiency of *nad7*-317 was increased in the mutants compared to ZH11, resulting in a higher rate of conversion of UCU (Ser) to UUU (Phe) (Figure **4A**). Furthermore, the editing efficiency of these 4 sites was also investigated in the complemented lines, and the results showed that the editing efficiency was restored along with the recovered expression levels of *PPR767* (Figure **4A**). The above results indicate that *PPR767* is related to the editing of 4 sites in 3 mitochondrial genes.

**Figure 4.**
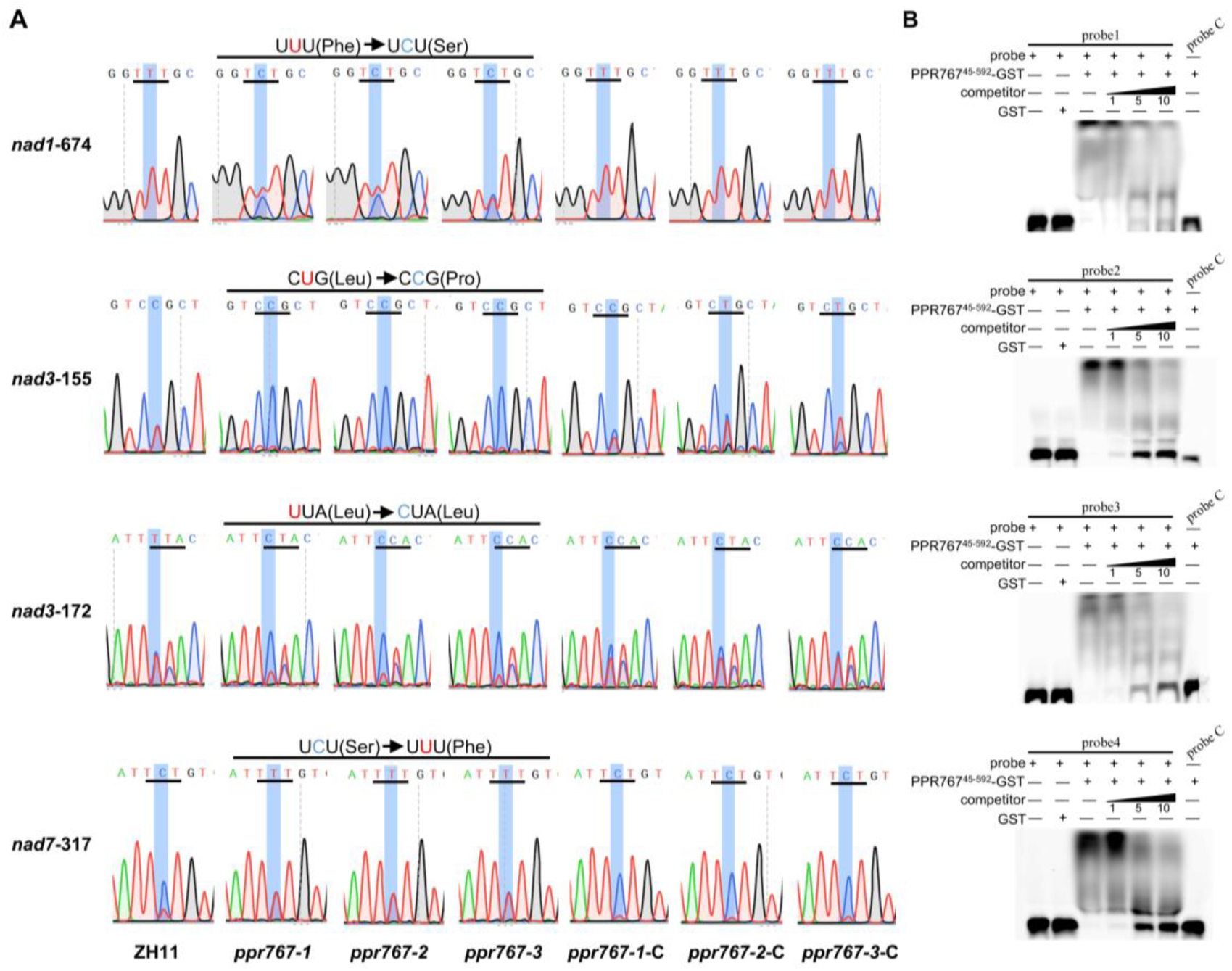
*PPR767* mutants alter the C-to-U editing efficiency of *nad1*, *nad3*, and *nad7* transcripts. **A**, The editing levels of mitochondrial genes at different sites: *nad1-674*, *nad3-155*, *nad3-172*, and *nad7-317*. The altered amino acids are indicated by black lines. **B**, Confirmation of the binding of PPR767 to five RNA targets by RNA electrophoretic mobility shift assays (REMSA). Unlabelled probes corresponding to the target probes were used as competitors. Probe C indicates negative control. Probes 1-4 indicate the corresponding probes with FAM tag to the target sites *nad1-674*, *nad3-155*, *nad3-172*, and *nad7-317*, respectively.

To investigate whether *PPR767* affects editing efficiency by directly binding to target RNA, we conducted RNA electrophoretic mobility shift assay (REMSA). In order to better express and purify the PPR767 protein in vitro, we excised 5’ and 3’ terminal sequences lacking functional domains or motifs, and the remaining sequence was amplified. The amplicons spanning 133 to 1776 bp (residues 45-592 aa) of *PPR767* were fused with GST tag protein sequence for expression. Furthermore, we synthesized approximately 35 bp RNA probes, which were labelled with FAM tag at 5’ terminus, along with corresponding unlabelled probes (cold probes) based on the target RNAs. As a negative control, the sequence of probe C was obtained from the nuclear genome. The specific sequences of the probes are listed in Supplemental Figure **S4**. Compared to the negative probe C, all the target RNA probe lanes of 4 editing sites emerged retarded bands after incubation with the PPR767^45-592^-GST fusion protein (Figure **4B**). Competitive experiments demonstrated that the binding shift intensity gradually decreased along with increasing concentrations of the corresponding competitive probes (Figure **4B**). These results indicate that PPR767 affects the editing efficiency of 4 mitochondrial editing sites by directly binding to target RNAs.

### PPR767 interacts with MORF1 and MORF8

As one element of the editosome, in addition to recognize and bind to target RNAs, PPR proteins also recruit other relevant proteins, including MORFs, to fulfil editing events (Takenaka et al., 2012). In this report, we screened MORF proteins that directly interacted with PPR767. Yeast two-hybrid (Y2H) assays revealed that only MORF1 and MORF8 interacted with PPR767 under the precondition that PPR767 lacked self-activation ability (Figure **5A**). To further verify this finding in vivo, a luciferase complementation imaging assay (LCI) was performed in tobacco leaves. Luciferase activity was detected in the sections in which MORF1 or MORF8 was cotransfected with PPR767 (Figure **5B-C**). These results were consistent with the results of the Y2H assay. Based on the above results, we conclude that both MORF1 and MORF8 interact with PPR767 and that PPR767 potentially recruits MORF1 and MORF8 to perform RNA editing.

**Figure 5.**
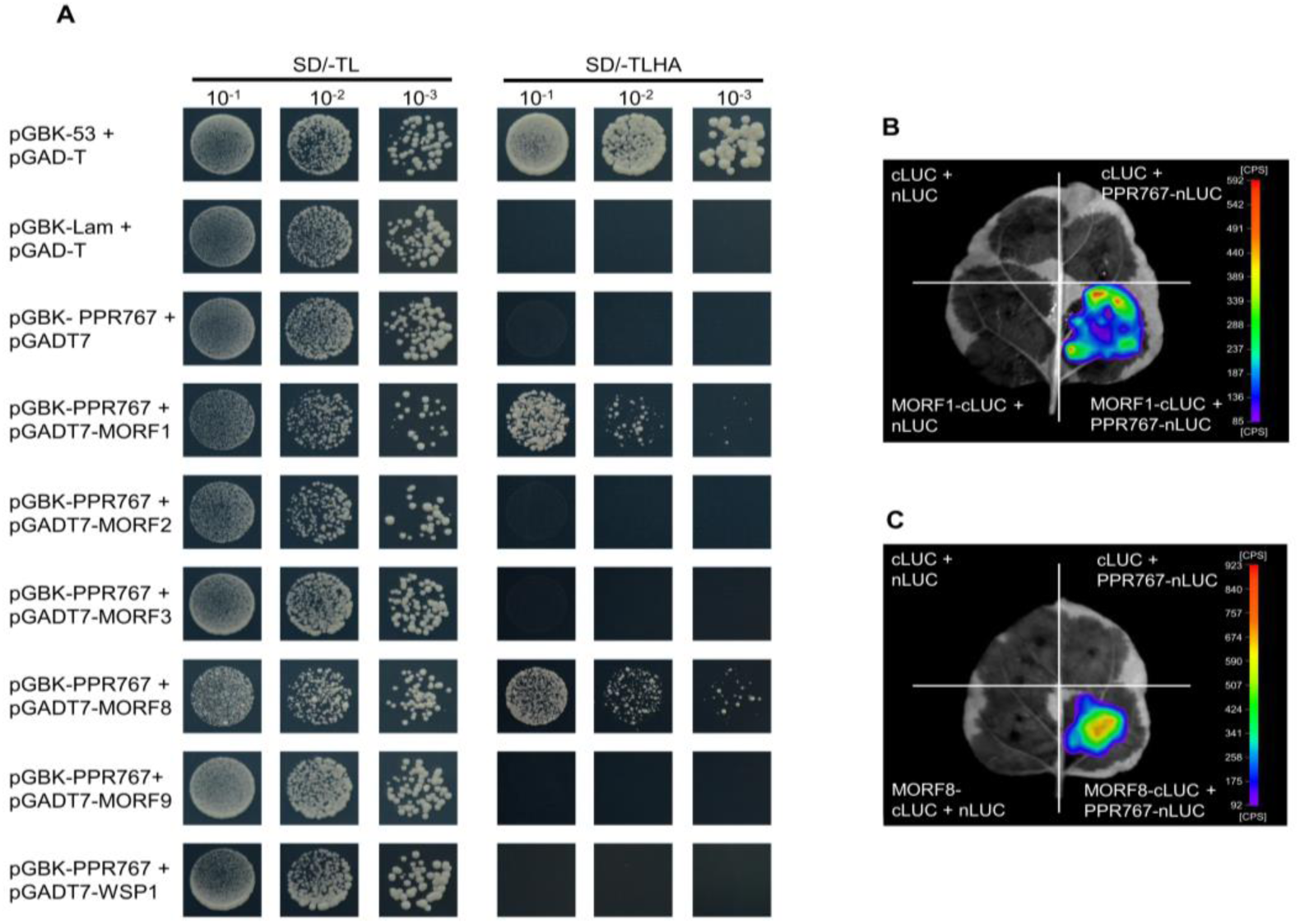
Protein interactions between PPR767 and MORFs. **A**, Yeast two-hybrid assays for interactions between PPR767 and MORF proteins. -TL and -TLHA indicate SD/-Trp-Leu and SD/-Trp-Leu-His-Ade medium, respectively. 10^-1^, 10^-2^, 10^-3^ indicate the dilution ratios of yeast cells. pGBK-53 and pGBK-Lam were utilized as positive and negative controls, respectively. pGADT7 was used for detection of self-activation. **B** to **C**, The verification of the interaction of PPR767 with MORF1 (**B**) or MORF8 (**C**) protein by LCI assay in tobacco leaves. cLUC and nLUC indicate the empty vector with the N-terminal or C-terminal domain of fluorescence, respectively.

Previous reports have demonstrated that the DYW-type PPR protein PPS1 is also involved in the editing events of *nad3*-155 and *nad3*-172 (Xiao et al., 2018b). Furthermore, PPR767, as an E-type PPR protein, lacks a DYW domain, indicating that it lacks the deaminase activity necessary for RNA editing events. Hence, we speculated that PPR767 may obtain deaminase activity from PPS1 to fulfil the editing events of *nad3*-155 and *nad3*-172. Initially, we investigated whether PPR767 directly binds to PPS1 via a Y2H assay. However, no yeast cells grew on the SD/-TLHA selection medium when PPR767-AD and PPS1-BD were cotransformed into Y2H gold cells (Supplemental Figure **S5A**). These results indicate that PPR767 does not directly interact with PPS1. Moreover, Y2H assay showed that MORF8 also interacted with PPS1 protein under the precondition that MORF8 lacked self-activation ability (Supplemental Figure **S5A**). According to previous results, both PPR767 and PPS1 interact with MORF8. To determine whether PPR767 indirectly interacts with PPS1 through MORF8 as a bridge, a yeast three-hybrid system was carried out. However, no yeast spots were observed on the medium, indicating that MORF8 does not serve as a bridge between PPR767 and PPS1 (Supplemental Figure **S5B**).

### Respiratory complex I shows reduced activity in *PPR767* mutants

The nad1, nad3 and nad7 subunits, which are encoded by mitochondrial genes, belong to the core subunits of complex I (Klusch et al., 2021). The correct modification of *nad1, nad3,* and *nad7* RNAs is crucial for complex I activity (Unseld et al., 1997). To investigate whether alterations about editing efficiency of the 4 editing sites caused by the mutation of *PPR767* affected the activity of complex I, we isolated mitochondria from calli of ZH11 and mutants and analysed them by blue native polyacrylamide gel electrophoresis (BN-PAGE). The purity of the mitochondria was confirmed by western blot (WB) assay. No RBCL or histone signals were detected in the mitochondria, indicating that they were pure and did not have contamination from the nucleus or chloroplast (Figure **6A**). The mitochondria freshly isolated from ZH11 and mutant calli were used to examine complex I activity. The results showed that the activity of complex I was significantly reduced in the mutant, indicating that the loss of function of PPR767 results in reduced complex I activity by influencing the editing efficiency of *nad1*-674, *nad3*-155, *nad3*-172, and *nad7*-317 (Figure **6B**).

**Figure 6.**
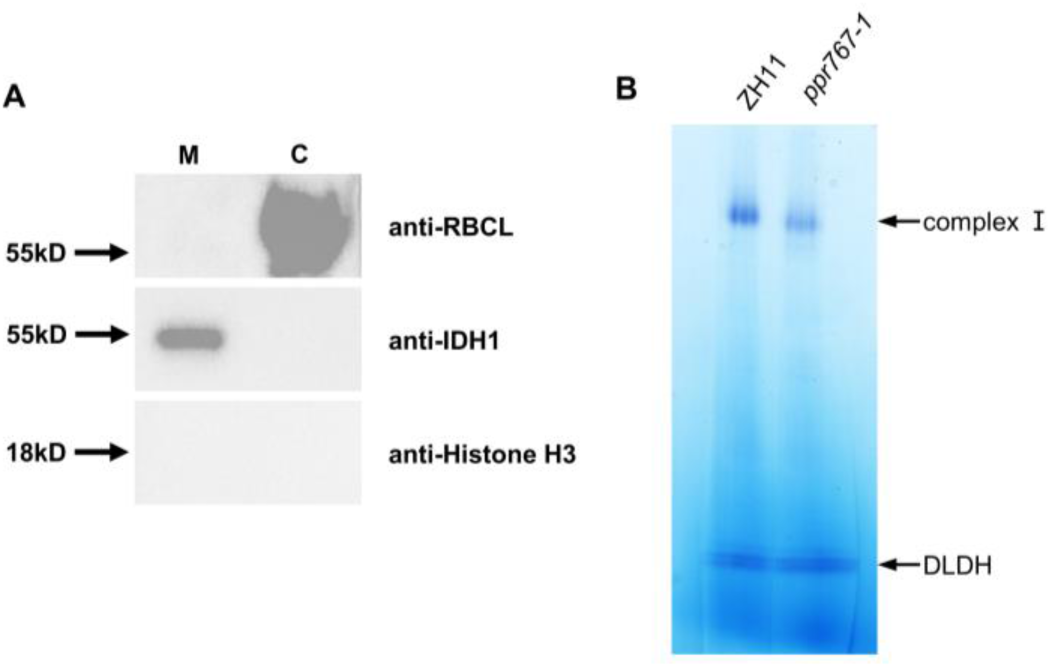
Mitochondrial complex I activity is decreased in mutants. **A**, Examination of mitochondrial purity. Mitochondria were extracted from calli. RBCL was used as chloroplast maker protein, IDH1 was used as mitochondria marker, Histone H3 was used as nuclear marker. M indicates isolated mitochondria, and C indicates purified chloroplast sample. **B**, Fresh mitochondria were extracted from the calli of ZH11 or *ppr767-1*. The mitochondrial complexes were separated by BN-PAGE. Arrows indicate the positions of complex I and dihydrolipoamide dehydrogemnase (DLDH), and the DLDH was used as a loading control.

### Mutation of *PPR767* increases ROS content and weakens drought resistance

Respiratory complex I, as the main entry point for electrons into the respiratory chain, affects the content of mitochondrial ROS. Previous reports have demonstrated that compromised complex I may cause an accumulation of ROS, which influences the stress resistance of plants (Moller, 2001; Lee et al., 2002; Zu et al., 2023). The reduced activity of complex I suggested that *PPR767* may affect stress resistance. Moreover, previous reports have demonstrated that the activity of complex I is related to plant immunity to pathogens, cold tolerance and so on, but the role of complex I in drought resistance is not well understood in rice (Yang et al., 2022b; Zu et al., 2023). In this study, we utilized two-week-old seedlings and performed RT-qPCR to analyse whether the expression level of *PPR767* could response to 15% PEG6000 treatment. The data showed that the expression of *PPR767* was induced in both the aboveground and underground parts and reached a peak after 8 h or 12 h of treatment, respectively (Supplemental Figure **S6**). These results indicate that *PPR767* is related to drought tolerance. To further confirm these results, we treated two-week-old seedlings of ZH11 and three mutants with 15% PEG6000, and compared to ZH11, fewer fresh leaves emerged in the mutants after recovery (Figure **7A**). And the statistical analysis verified that the survival rate of the WT was significantly higher than that of the mutants after recovery (Figure **7B**). These results reveal that the compromised activity of complex I lead to reduced drought resistance in *PPR767* mutants.

**Figure 7.**
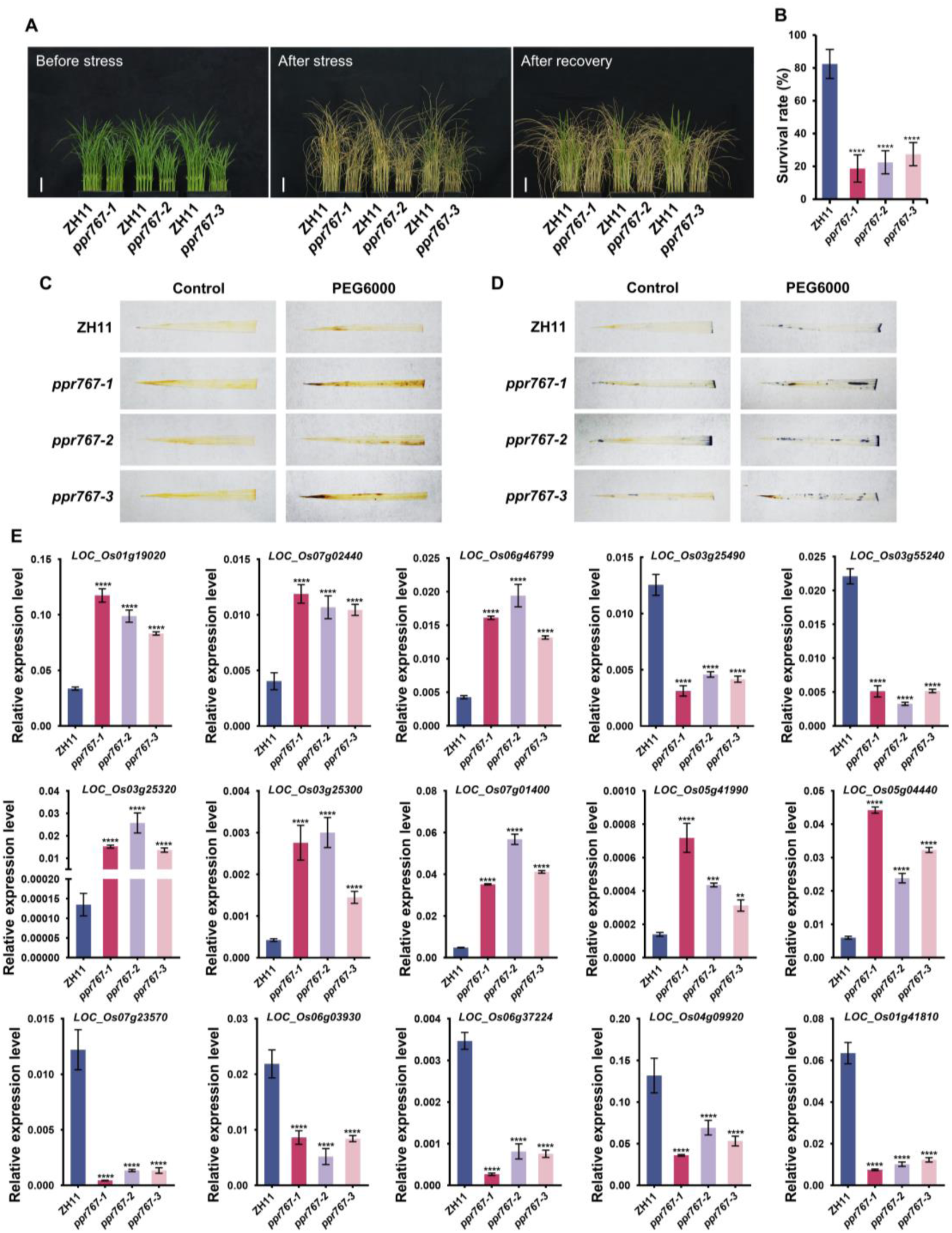
*PPR767* positively regulates drought tolerance. **A**, Phenotype of ZH11 and mutants seedlings (cultured for two weeks in black bottles after sowing) after drought stress and after recovery. Scale bar =5 cm. **B**, Statistical analysis of survival rate of ZH11 and the mutants after recovery. Data are means ± SDs of three biological replicates. **C** to **D**, DAB **(C)** and NBT **(D)** staining of ZH11 and mutants leaves. The leaves were collected from two-weeks-old seedlings treated with or without 15% PEG6000 treatment for 8 h. Control and PEG6000 indicate before and after stress treatment, respectively. **E**, RT-qPCR assay to test the expression levels of differentially expressed genes (DEGs). Data are means ± SDs of biological triplicates. Asterisks indicate significant differences by Student’s two-tailed *t* test (*\*p* < 0.05, *\*\*p* < 0.01, *\*\*\*p* < 0.005, *\*\*\*\*p* < 0.001, ns, no significance).

In order to investigate whether *PPR767* affected drought tolerance by influencing the ROS content, we performed 3,3’-diaminobenzidine (DAB) and nitroblue tetrazolium (NBT) staining with leaves of two-week-old seedlings to visualize the content of hydrogen peroxide (H_2_O_2_) and superoxide anion radical (O_2_−), respectively. Compared to the WT, a larger portion of the leaves was stained by NBT or DAB in the mutants after 24 h treatment with 15% PEG6000, indicating that the knockout of *PPR767* leads to the accumulation of ROS under drought treatment (Figure **7C-D**). Meanwhile, two-week-old seedlings of ZH11 or *ppr767-1* mutants were separated into aboveground and underground parts for transcriptome analysis. In the aboveground samples, 619 differentially expressed genes (DEGs) were identified, including 359 upregulated genes and 260 downregulated genes in the mutant (Supplemental Figure **S7A-B**). Gene Ontology (GO) analysis revealed that several DEGs were enriched in the ‘oxidation-reduction process’ term (Supplemental Figure **S7C**). In underground samples, 2035 DEGs were identified, with 1226 upregulated DEGs and 809 downregulated DEGs in *ppr767-1* (Supplemental Figure **S8A-B**). Several DEGs were enriched in ‘response to oxidative stress’, ‘peroxidase activity’, ‘oxidoreductase activity, acting on paired donors, with incorporation or reduction of molecular oxygen’, and ‘oxidoreductase activity, acting on single donors with incorporation of molecular oxygen, incorporation of two atoms of oxygen’ GO terms (Supplemental Figure **S8C**). Upon detailed analysis of these DEGs, we found that several genes annotated by the UniProt website (https://www.uniprot.org/) were associated with the production or scavenging of ROS. And most up-regulated genes were annotated as peroxidase which is involved in the elimination of ROS, while most of the downregulated genes were annotated to the cytochrome P450 superfamily, which is involved in the induction of ROS production in both aboveground and underground samples. To verify the RNA-seq results, we selected several genes associated with ROS for RT-qPCR analysis, including the significantly upregulated genes *LOC_Os01g19020*, *LOC_Os07g02440*, and *LOC_Os06g46799*, and the significantly downregulated genes *LOC_Os03g25490*, and *LOC_Os03g55240* in aboveground samples; the significantly upregulated genes *LOC_Os03g25320*, *LOC_Os03g25300*, *LOC_Os07g01400*, *LOC_Os05g41990*, and *LOC_Os05g04440*, and the significantly downregulated genes *LOC_Os07g23570*, *LOC_Os06g03930*, *LOC_Os06g37224*, *LOC_Os04g09920*, and *LOC_Os01g41810* in underground parts of the mutants. The expression patterns of these genes, as determined by RT-qPCR, were consistent with the RNA-seq data (Figure **7E**). These results indicate that the mutation of *PPR767* affects ROS accumulation. In summary, the above results demonstrate that the reduced complex I activity caused by mutation of *PPR767* results in increased ROS content, which impairs the drought resistance of the mutants.

### Predicted alignment of PPR767 to the RNA editing target sequences

The recognition model of the PPR protein, which is recognized as RNA-binding protein, has been predicted in some reports (Barkan et al., 2012; Takenaka et al., 2013). The model indicates that the amino acids at positions 1’ and 6’ of motifs are crucial for recognizing the upstream sequences of target RNA sites. The amino acid at position 6’ refers to the sixth amino acid within a given PPR motif, while the amino acid at the 1’ position refers to the first amino acid of the subsequent adjacent motif at the C-terminus. The fourth upstream nucleotide from the editing site is recognized by the 1’ amino acid of the E motif and the 6’ amino acid of the adjacent S2 motif. The N-terminal motifs preceding the S2 motif sequentially recognize the upstream sequences of the forth nucleotide (Takenaka et al., 2013). Based on previous reports, we inferred the standard recognition sequences of PPR767 and aligned them with the upstream nucleotides of *nad1*-674, *nad3*-155, *nad3*-172, and *nad7*-317, respectively. Because PPR767 contains 13 PLS motifs, the corresponding recognition upstream sequences from the 4th to the 16th nucleotides of the target sites were listed in the alignment (Figure **8A**). We also calculated the predicted nucleotide binding intensities (predicted matches/all matches) for each PPR repeat in the PPR767 protein (Figure **8B**). A higher predicted score indicates stronger binding intensity. The results showed that the PPR motif of PPR767 was more closely related to the target sequence.

**Figure 8.**
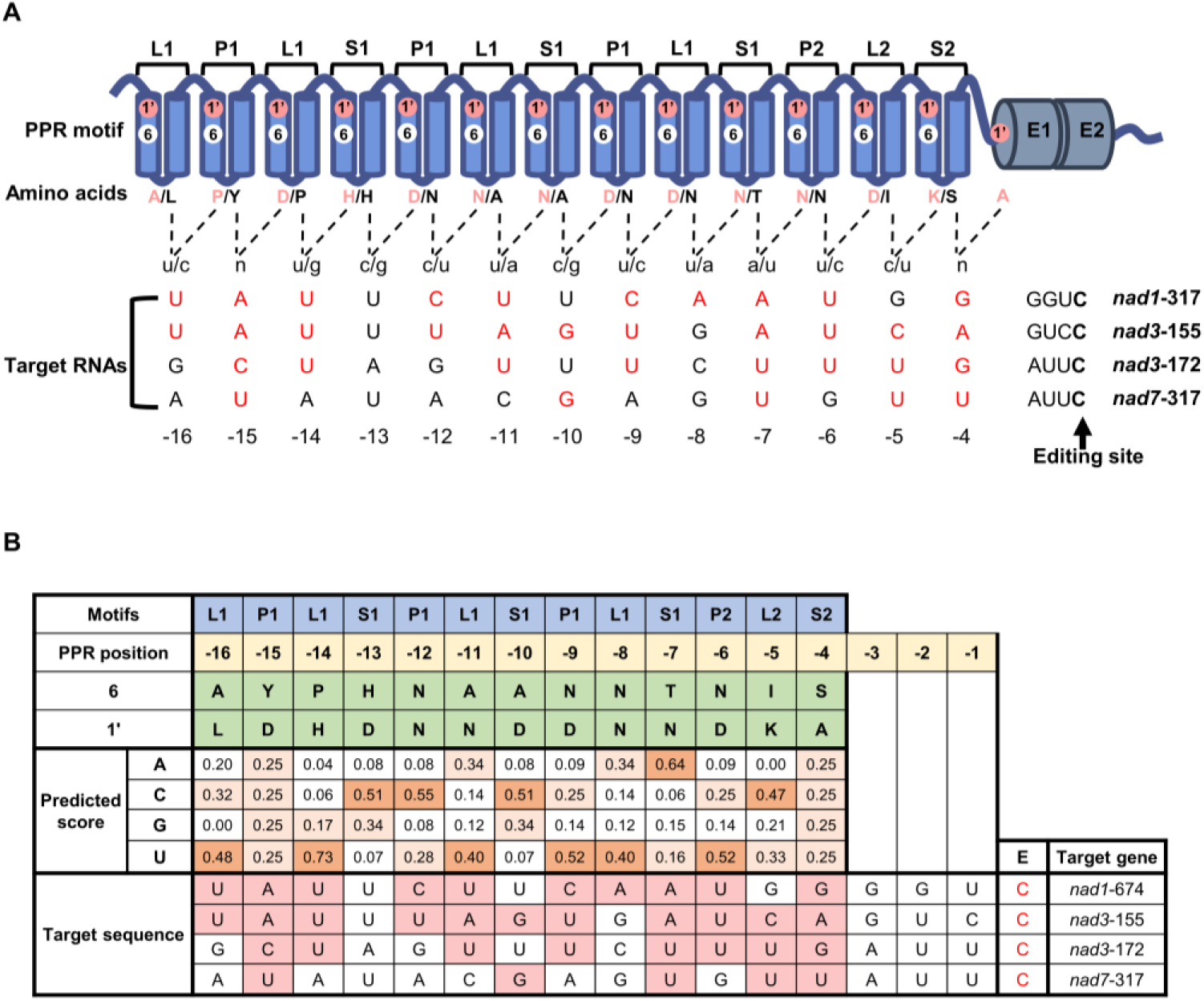
The alignment and prediction of PPR motifs contained by PPR767 to the target RNA editing sequence. **A**, The alignment of motifs to target RNA sequences. Position 1’, indicated by rosy circle, and position 6, indicated by white circle, indicate the first and sixth amino acids of correlated PPR motif, respectively. The corresponding amino acids are listed under the motif structure and labelled with identical color. The dashed lines and the lowercase letters of nucleotides indicate the presumed connection to target nucleotide identities. The bold “C” pointed by arrow indicates the editing site. The uppercase letters indicate the upstream sequence of the correlated editing sites, and the red letters indicate the nucleotides that fit the recognition rule. The number indicates the specific order of nucleotides. **B**, Prediction of the target RNA sequence of PPR767. The PPR motifs are shown in blue part and their alignment to nucleotide positions which are counted 3’ to 5’ from the edited C (from right to left, −4 to −16) is labelled by yellow background. Amino acids at position 6 (the sixth amino acid of the considered motif) and 1’ (the first amino acid of the next C-terminal PPR motif) are marked by green, and the corresponding predicted score of every site is shown in the below table cell (more intense orange color denotes higher score). The next several lines show the known target sequences, and the target nucleotides corresponding to the recognition model are marked by rose red. The red font “C” indicates editing sites.

## Discussion

In flowering plants, mitochondria as organelles to supply energy, the accurate expression of mtDNAs is important and requires the involvement of nucleus-encoded proteins (Knoop, 2004; Barkan and Small, 2014). PPRs, as nucleus-encoded RBPs, participate in mitochondrial RNA metabolism by directly binding target RNAs and impact plant development and stress tolerance (Yin et al., 2013). Several previous reports have revealed that loss of PPRs function impacts cold resistance, immunity to pathogens, and salt tolerance by influencing mitochondria complex activity and ROS content in rice. Nevertheless, it is not well elucidated that the function of RNA processing of respiratory complexes in drought resistance in rice.

In this study, for stem transverse sections, we did not measure the interval of ABVs in the fourth internode because the alignment of ADVs in the fourth internode is dramatically irregular. For leaf transections, the intervals of LVs were not measured, because they are determined by the number and interval of SVs, and the relevant data are shown in this report.

In this report, we discovered that PPR767 participates in the editing events of *nad3*-155, and *nad3*-172, and coincidentally, PPS1, which has been studied, is related to these events as well (Xiao et al., 2018b). However, based on these results, PPR767 does not interact with PPS1 either directly or indirectly via MORF8 as a bridge (Supplemental Figure **S5A-B**). According to the results shown in Figure **4A**, compared to WT, the editing efficiency of *nad3*-155 and *nad3*-172 in the *ppr767* mutants was partially decreased. This phenomenon likely demonstrates that more than one PPR protein is involved in the editing of the same editing site. Notably, although both PPS1 and PPR767 can bind to target RNA directly, the editing efficiency of these two sites drops to zero when *PPS1* is knocked out. The complete interruption of the editing event in *pps1* mutants demonstrates that PPS1 is significant and indispensable for these editing events. This result also suggests that the editing complex raised by PPR767 should include PPS1 in these two editing events and that other yet-to-be-identified factors that serve as a bridge between PPS1 and PPR767 must be recruited by PPR767 to obtain deaminase activity. Hence, the elements in the complex raised by PPR767 need to be further investigated.

The editing efficiency of *nad7*-317 is increased in the *PPR767* mutants (Figure **4A**). It is hypothesized that the redundancy about the cis-acting elements involved in the editing of *nad7*-317, suggesting that additional PPR proteins participate in this editing event. And this phenomenon may indicate that PPR767 also inhibits the binding of other PPR proteins to this target RNA by occupying the upstream sequences of the editing site.

Mutation of *PPR767* causes alters editing efficiency of 4 editing sites in 3 mitochondrial genes and reduces the activity of complex I, resulting in increased ROS content, and decreased drought resistance. These findings indicate that electron transport in complex I has an essential effect on the ROS content. This finding is similar to that of a previous report about Arabidopsis *fro1* mutants, in which these mutants also give rise to a deficiency of complex I, an ROS burst, and subsequently the reduced cold resistance (Lee et al., 2002). Furthermore, in our study, RNA-seq revealed that the expression levels of nucleus encoded genes, some of which are related to ROS production or elimination, are altered in the mutants due to the compromised function of complex I. These phenomena also provide references and evidence for retrograde signals from mitochondria to the nucleus.

Furthermore, it has been reported that RNA processing in mitochondria could be involved in stress resistance by affecting complex I activity. The *Arabidopsis* protein AtRTP7 functions in both RNA splicing of mitochondrial gene *nad7* and RNA editing of *nad1*, *nad3*, and *nad6* and further reduced the activity of complex I. These mutants result in mROS burst and enhanced resistance to plant disease (Yang et al., 2022b). The rice protein SOP10 has been reported to function in RNA splicing of *nad4* and *nad5* transcripts and affect resistance to cold stress by reducing NADH dehydrogenase activity (Zu et al., 2023). However, the mechanism by which PPRs influence osmotic stress resistance via impacting complex I activity in rice has not been fully elucidated. In this work, we show that the knockout of *PPR767*, which functions in the RNA editing of *nad1*, *nad3*, and *nad7* transcripts, causes reduced complex I activity and further leading to an increase in ROS level and a decrease in drought tolerance in *japonica* rice, providing a potential target for breeding of drought resistance in *japonica* rice.

In summary, we identified the E-type PPR protein PPR767 is an RNA processing factor of complex I subunit transcripts, and the loss of PPR767 functionality lead to growth reduction, ROS burst, and reduced drought tolerance. These results indicate that numerous PPRs may play important roles in plant drought resistance.

## Materials and methods

### Plant materials

The wide type *japonica* rice (*Oryza sativa*) cv “Zhonghua11” was used as the background for the transgenic lines. The *PPR767* KO lines generated by the CRISPR/Cas9 system were purchased from BIOGLE GeneTech Company (Hangzhou, China). Both the WT and transgenic plants were grown in paddy fields under natural conditions with proper management.

For seedling analyses, the germinated seeds were sowed in the black and opaque box with a 96-well bottomless black PCR plate. The plants were grown at 28℃ in a growth room, with a 10-h-light/14-h-dark cycle and cultured with Yoshita liquid medium (Coolaber, China). Calli derived from the WT and the mutants were cultured on Murashige and Skoog subculture medium.

### Phenotype analysis and stress treatment

Field phenotypes including plant height, flag leaf length and width, internode length and thickness, tiller number, grain number, as well as number of primary and second branches were measured at the mature stage. For drought treatment, two-week-old seedlings were treated with Yoshita liquid medium containing 15% (w/v) PEG6000 for 14 days. After recovery, the seedlings with new fresh leaves were counted as the survivors.

### Plasmid construction and plant transformation

The amplicons of the whole genome sequence of *PPR767*, containing the 2000 bp sequence upstream of the 5’ end of *PPR767*, were amplified and subcloned and inserted into pCAMBIA2300 vector. The construct was introduced into *A. tumefaciens* strain GV3101, which was subsequently transformed into the knock out lines which were homozygous and lack Cas9 element, and the complemented lines were obtained. Primers used in this study are listed in supplemental Table **S1.**

### Subcellular localization of PPR767

The CDS in which the terminator codon of *PPR767* was removed was amplified from the reverse transcription product of ZH11 mRNA. Amplicons were subsequently cloned and inserted into the HBT-sGFP vector. The fusion protein of PPR767 and green fluorescence protein was driven by the cauliflower mosaic virus (CaMV) 35S promoter. The constructed vector was introduced into rice protoplasts using a polyethylene glycol-mediated transformation system as previously described (Yang et al., 2022a). Confocal laser scanning microscope (Leica Microsystems, Germany) was employed to acquire fluorescence of 35S::PPR767-GFP. The MitoTracker Red (YEASEN, China) was used as mitochondrial indicator. Primers used in this study are listed in supplemental Table **S1.**

### Measurement of the net photosynthetic rate

The net photosynthetic rate was measured by LI-6800 system (LI-COR, USA). The middle of the flag leaves at seven days after heading was used for the measurements. The concrete methods were described previously (Fan et al., 2023).

### Histological analysis

For histological analysis, the end of each internode and the middle segment of flag leaves were fixed in FAA solution (3.8% [v/v] formaldehyde, 5% [v/v] acetic acid, and 63% [v/v] ethanol). After fixation for more than 24 h, the tissues were embedded in paraffin. The paraffin slides were dewaxed and dyed with safranin O staining solution and plant solid green staining solution. The prepared sections were observed under stereomicroscope SMZ25 (Nikon, Japan) and images were taken for analysis. The size and interval of vascular bundles were analysed and measured with ImageJ software (https://imagej.nih.gov/ij/).

### Transmission electron microscopy assays

For the TEM assay, the leaves of two-week-old seedlings were cut into small pieces of approximately 2 × 2 cm^2^, and fixed rapidly in 0.1 M phosphate buffer (pH 7.4) supplemented with 2% (v/v) glutaraldehyde and chilled on ice. Then, the samples were kept in vacuum untill the pieces sank into solution. The methods used were described previously (Xiao et al., 2018a). The samples were observed with transmission electron microscope (JEM-1400plus, Japan).

### RNA extraction and RT-qPCR

Total RNA was extracted from the tissues using TRIzol Reagent (Thermo Fisher, USA). First strand cDNA was synthesized with a Maxima H Minus First Strand cDNA Synthesis Kit with dsDNase (Thermo Fisher, USA) following the kit instructions. RT-qPCR was performed using SYBR Green PCR mix (YEASEN, China) on a LightCycler 480 system (Roche, Switzerland). The rice *Actin* gene was used as the endogenous control for normalization. The primers used for analysis are listed in Table S1. The PCR program was performed according to the manufacture’s instruction.

### RNA editing

Total RNA was extracted from leaves of the WT and transgenic plants. cDNAs were synthesized via reverse transcription reactions with random primers. In order to certify the editing sites, primers covering all known mitochondrial and chloroplast editing sites were designed and listed in Supplemental Table **S1**. The Sanger sequencing results of the PCR products were utilized to analyse the editing efficiency.

### RNA electrophoretic mobility shift assay (REMSA)

The corresponding cDNA fragments of *PPR767* were amplified, and the product was cloned and inserted into pGEX-6P-1 vector. The constructed vector was subsequently transformed into *E. coli.* strain BL21. The fusion protein PPR767^44-592^-GST was expressed in vitro and subsequently purified by GST-tag beads (YEASEN, China). The RNA probes and negative control probe were labelled with FAM, and these labeled probes and competing probes were synthesized by GeneScript (Nanjing, China). The REMSA was performed as described previously (Xiao et al., 2018b). The polyacrylamide gel was imaged by laser scanner (Typhoon, USA).

### Yeast two-hybrid (Y2H) assay

The full-length CDSs of *PPR767* and *PPS1* were cloned and inserted into pGBKT7 vector respectively, while the CDSs of *PPR767*, *MORF1*, *MORF2*, *MORF3*, *MORF8*, *MORF9* and *WSP1* were cloned and inserted into pGADT7 respectively. The constructed vectors were transformed into the yeast strain Y2Hgold. The transformed yeast cells were inoculated on SD/-Leu/-Trp medium for 48-72 h at 29 ℃. The diluted yeast cells were transferred to new SD/-Leu/-Trp, SD/-Leu/-Trp/-His and SD/-Leu/-Trp/-His/-Ade medium. Primers used in this study are listed in supplemental Table **S1.**

### Luciferase complementation imaging assay (LCI)

For the LCI assay, the CDS without termination codon of PPR767 was amplified and cloned and inserted into the nLUC vector. The constructed vector expressed the PPR767-nLUC fusion protein. The CDSs without stop codon of MORF1 and MORF8 were linked to the cLUC vector respectively to generate the MORF1-cLUC and MORF8-nLUC fusion proteins respectively. The experiment was performed following previous reports (Chen et al., 2024). Primers used in this study are listed in supplemental Table **S1.**

### Yeast there-hybrid (Y3H) assay

For the Y3H assay, the coding sequence of PPS1 was cloned and inserted into multiple cloning site I (MCS I) of the pBridge vector. The CDS of MORF8 protein was cloned and inserted into MCS Ⅱ. The CDS of PPR767 was cloned and inserted into pGADT7 vector. All the constructed vectors were transformed into the yeast strain AH109. The transformed yeast cells were inoculated on SD/-Leu/-Trp/Met medium for 48-72 h at 29℃. The diluted yeast cells were transferred to new SD/-Leu/-Trp/Met, and SD/-Leu/-Trp/-His/-Met medium for 48-72 h at 29℃. Primers used in this study are listed in supplemental Table **S1.**

### Blue native polyacrylamide gel electrophoresis (BN-PAGE)

Mitochondria were extracted from calli, and mitochondrial proteins were extracted following previous methods (Yang et al., 2022a). The BN-PAGE and assessment of complex I activity were performed as described previously (Xiao et al., 2018b). The gel was scanned with a scanner (Epson Perfection V700 Photo, Japan).

### Detection of ROS

The leaves of two-week-old seedlings treated with 15% PEG6000 for 24 h or without treatment were the samples for histochemical staining. The 3,3’-diaminobenzidine (DAB) and nitroblue tetrazolium (NBT) staining were carried out according to corresponding instructions (Servicebio, China).

### RNA-seq

RNA samples (three biological replicates per condition) were sent to company for sequencing. RNA-seq experiment and high throughput sequencing were conducted by Seqhealth Technology Co., Ltd. (Wuhan, China). 2 μg total RNA was used for stranded RNA sequencing library preparation. The de-duplicated consensus sequences were mapped to the reference genome of MSU7.0 (version 2.5.3a) with default parameters. For data analysis, a p-value cut-off of 0.05 and fold-change cut-off of 2 were used to judge the statistical significance of the differences in gene expression. Gene Ontology (GO) enrichment analysis of differentially expressed genes was implemented with g:Profiler (https://biit.cs.ut.ee/gprofiler/gost).

### Accession numbers

Sequence information can be found in the Rice Genome Annotation Project (http://rice.plantbiology.msu.edu/) under the following accession numbers: *LOC_Os05g24150* (*PPR767*), *LOC_Os11g11020* (*OsMORF1*), *LOC_Os06g02600* (*OsMORF2*), *LOC_Os03g38490* (*OsMORF3*), *LOC_Os09g33480* (*OsMORF8*), *LOC_Os08g04450* (*OsMORF9*), *LOC_Os04g51280* (*WSP1*), *LOC_Os12g36620* (*PPS1*), *LOC_Os01g19020*, *LOC_Os07g02440*, *LOC_Os06g46799*, *LOC_Os03g25490*, *LOC_Os03g55240*, *LOC_Os03g25320*, *LOC_Os03g25300*, *LOC_Os07g01400*, *LOC_Os05g41990*, *LOC_Os05g04440*, *LOC_Os07g23570*, *LOC_Os06g03930*, *LOC_Os06g37224*, *LOC_Os04g09920*, *LOC_Os01g41810*.

## Acknowledgements

We thank Yukun Wang (Wuhan University) for kindly providing the protein sample of pure chloroplast. We thank Yunping Chen, Zhiwu Dan, Fengfeng Fan, and Manman Liu from Wuhan University for providing assistance in the experimental technology. This work is supported by the National Natural Science Foundation of China (31871592), the Creative Research Groups of the Natural Science Foundation of Hubei Province (2020CFA009), and the Fundamental Research Funds for the Central Universities (2042022kf0015). We thank Dr. Wei Huang from Guangzhou biomedshine Co.Ltd is highly appreciated for his donation to this project.

## Author contributions

LP and JH designed the research. LP performed most of the experiments. HX, YX and XY performed additional experiments. ZH, CL and LH collected the data about yeild trait and contributed to field management. LP and JH wrote the manuscript.

